# Relationship between ocean area and incidence of anthropogenic debris ingested by longnose lancetfish (*Alepisaurus ferox*)

**DOI:** 10.1101/578310

**Authors:** Mao Kuroda, Keichi Uchida, Yoshinori Miyamoto, Ryuiti Hagita, Daisuke Shiode, Hiroki Joshima, Masaki Nemoto, Hiroaki Hamada, Yuta Yamada, Hideshige Takada, Rei Yamashita, Kohei Fukunaga

## Abstract

Longnose lancetfish (*Alepisaurus ferox*) may has been studied as an indicator of marine pollution caused by marine litter. The objectives of this study were to determine the difference in frequency of occurrence of plastics ingested by longnose lancetfish in different ocean area. In this study, we compared the incidence and characteristics of anthropogenic debris in the stomachs of longnose lancetfish. We examined 91 longnose lancetfish caught by pelagic longline fishing in Sagami Bay, the North Pacific Ocean, approximately 200 km south of Shikoku, and in the Indian Ocean. Broken down by ocean area, the incidence of anthropogenic debris ingestion was highest in Sagami Bay (23 of 34 specimens, 68%), followed by the North Pacific Ocean (1 of 9, 11%), and the Indian Ocean (8 of 48, 17%). The frequency of occurrence increased in area close to the sphere of human habitation. The anthropogenic debris collected in this study were more than 70% classified as plastic sheeting. Stomach content analysis revealed that more than 90% of the plastic fragments were composed of PP and PE, which have specific gravities that are less than that of seawater. The results of this study show that some of the plastics flowing from the land into the sea are spreading through under the water surface of the ocean.

## Introduction

The issue of marine debris has received considerable interest in recent years. Marine debris is studied mainly into three types: shoreline debris that washes ashore[1],[2], floating debris that floats on the ocean surface[3],[4], and ocean floor debris that is deposited on the ocean floor[5],[6],[7]. Most marine debris is composed of plastic, primarily because it is light strong and durable and well suited for becoming marine debris; i.e., it floats on water and can be transported long distances without being degraded [8]. Plastic debris in the ocean can be ingested by marine animals, or they can become ensnared in it, can cause harm to marine mammal[9],[10]. Over time, plastic debris is broken down into increasingly small fragments by UV light, temperature changes, and wave action. Plastic fragments measuring 5 mm or less in size that are created by these processes are referred to as microplastics. Microplastic also includes plastic less than 5 mm from original, such as microbeads and resin pellets. Although it has been confirmed that plastics will be reduced to nano size, they never disappear and remain in the environment for an extremely long time. Furthermore, such microplastic fragments act as a medium that can concentrate persistent organic pollutants in seawater, raising concerns that their ingestion will promote the uptake of persistent organic pollutants by organisms [11].

Considerable research has been conducted on the extent and status of marine debris and its impact on marine life to date. A large proportion of this research has examined the effects of marine debris on seabirds. In a study conducted in 1997, plastic was detected in 97.6% of juvenile birds sampled [12]. According to Wilcoxa et al. (2015), if this trend continues, plastic fragments will likely be found in all seabird species by 2050 [13]. In this way, researching the relationship between seabirds and plastic fragments contributes to prediction of the increase in microplastic.

On the other hand, some studies have evaluated the change in dust in the area by investigating the stomach contents of the fish for a long time. Among fish species, research on the deep-water longnose lancetfish (Alepisaurus ferox, Lowe 1833) has a relatively long history.

In 1964, plastics were found in the stomach of longnose lancetfish in Suruga Bay of Shizuoka prefecture in Japan [14]. Longnose lancetfish (Alepisaurus ferox Lowe) is widely distributed in the oceans of the worlds of the Pacific, Atlantic and Indian Oceans excluding bipolar regions. The body length will be around 2 m, when they grow. This species belongs to the genus longnose lancetfish, contains 2 genera in this department, one species (A. brevirostris Gibbs) does not live in the North Pacific region. Both species are deep-sea fish and there is no fisheries value because muscle contains a lot of water and it is inedible[15]. This species is among the most common bycatch species from tuna longline fishing [16]. Also, longnose lancetfish has the characteristic of Stomach contents are generally well preserved because food is stored in the stomach and digested in the intestines [17].

The wide variation of sizes, textures, colors, and shapes of stomach contents demonstrate the opportunistic feeding behavior and lack of selectivity [18], and It is known that reflects the composition of the animal society to which longnose lancetfish belongs in each ocean area [14]. From these characteristics and research results that longnose lancetfish is suitable for use as an indicator species of marine pollution, longnose lancetfish has been regarded as an indicator species to examine the actual condition of marine debris.

However, although it has been reported that longnose lancetfish is an indicator species for marine debris, there are few cases where the results are compared between waters. In previous studies, many surface drifting marine debris observations have been conducted. Although marine debris needs to be examined, not only at the ocean surface, but also in the water column. Therefore, in this study, we compared the incidence and characteristics of artificial debris in the stomach of longnose lancetfish collected in the coastal areas of Japan, the Pacific Ocean, and the Indian Ocean. Based on the results, we tried to compare the actual condition of marine debris between ocean area.

## Materials and Methods

### Sample collection

We examined 91 longnose lancetfish caught by longline fishing during 22 fishing operations in Sagami Bay aboard the Seiyo-maru: 170 GT from October 2013 to February 2017; three fishing operations in the North Pacific Ocean, approximately 200 km south of Shikoku aboard the Shinyo-maru: 694 GT in September, 2015, and 21 fishing operations in the Indian Ocean aboard the Umitaka-maru: 3391 GT from December, 2014 to December, 2015 to December, 2016 (Fig. 1). In all cases, fishing was conducted during the daytime, with fishing lines set in the morning and then reeled in later the same day. Thawed frozen mackerel with fork lengths of 20 to 27 cm were used as bait. Of the specimens caught, 34 were caught in Sagami Bay, 9 were caught in the North Pacific Ocean, and 48 were caught in the Indian Ocean. All specimens were stored in a freezer immediately after caught and stored frozen. And after the voyage, we analyzed them in the laboratory. At first the frozen specimens were thawed that standard length (cm) and wet weight (g) were measured. The stomach was excised similar to Jantz et al (2013)[19]. The excised stomach contents were treated similar to jackson et al (2000)[20]. Fish were not sorted by sex because A. ferox are synchronous hermaphrodites, where the ovarian and testicular tissues are simultaneously developed [21]. Plastic marine debris pieces <1 mm in size were not quantified in this study as they were difficult to see with the naked eye [22]. However, in this study, we couldn’t find anthropogenic items of micro size (< 5 mm). For this reason, we targeted anthropogenic items of macro size (> 5 mm). In addition, the capture depth of each specimen was estimated based on the design of the longline fishing gear and data from depth-loggers attached to some of the fishing gear. The capture depth was not estimated for 12 specimens that had no record of hook number.

**Fig. 1.**
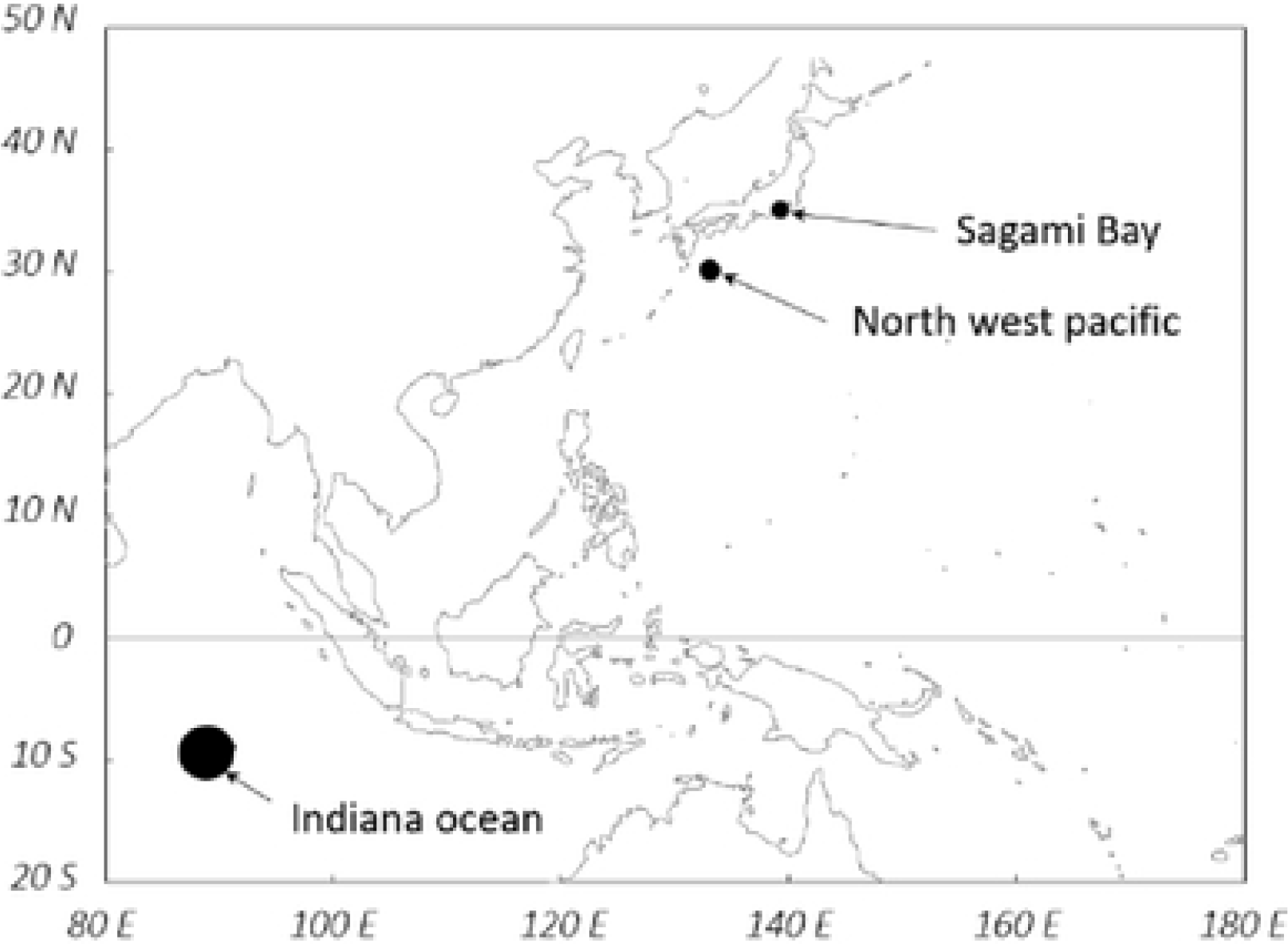
Map of sample areas: 26 km off the coast in Sagami Bay, 285 km (southward) off the coast of Shikoku in the northwestern Pacific, 1660 km off the coast in the Indian Ocean.

### Stomach content analysis

After separating the extracted stomach contents into anthropogenic and natural items, the anthropogenic items were further separated by shape and feel referred to the classification scheme of Jantz et al (2013)[19]. Briefly, the items were sorted into seven items: plastic sheet, plastic piece, rope piece, miscellaneous items, paper, rubber, yarn.

After that, the plastic sheet was sorted into plastic bags and food packaging items by printing, color and feel. We then measured the longest axes, area, dry weight (g) of the anthropogenic and natural items that were obtained. The stomach contents of 14 specimens caught in Sagami Bay, 8 specimen caught in the Indian Ocean, and one specimen caught in the North Pacific Ocean, were photographed, and the projected areas and longest axes of different items were calculated using a Microsoft Excel macro “!0_0! Excel Length and area measurement” (http://www.vector.co.jp/soft/win95/art/se312811.html). Weight(g) was measured using an analytical scale up to 0.1g.In addition, the anthropogenic stomach contents of ten specimens caught in Sagami Bay, eight specimen caught in the Indian Ocean, and one specimen caught in the North Pacific Ocean, were analyzed by Fourier transform infrared (FT-IR) spectroscopy (Nicolet iS5, Thermo Scientific) and PlaScan-W (Systematic engineering Inc. Japan), the materials were specified.

### statistical analyzes

All statistical analyzes were performed using Microsoft Excel for office 365 MSO. Correlation between the length of longnose lancetfish and the size of anthropogenic items ingested, standard length and capture depth, correlation between the number of anthropogenic items and the capture depth was obtained using the scatter chart. The chi-square test was used to confirm the significance of the comparison of results for each ocean area. The significance level was set to α = 0.05.

## Results

### Incidence of anthropogenic debris ingestion

Anthropogenic items were found in the stomach contents of 32 out of 91 specimens, or 35% of the specimens. Broken down by ocean area, the incidence of anthropogenic debris ingestion was highest in Sagami Bay (23 of 34 specimens, 68%), followed by the Indian Ocean (8 of 48, 17%), and the North Pacific Ocean (1 of 9, 11%)(Fig. 2a). We then examined the relationship between body length and the incidence of anthropogenic debris ingestion. As shown in Fig. 2b, the incidence of ingested plastics was markedly lower in smaller specimens. From the body length distribution in different ocean areas (Fig. 2c), small specimens were common in the Indian Ocean, where there was low incidence of anthropogenic debris, and in the North Pacific Ocean. These results suggest that the difference in the incidence of anthropogenic debris ingestion could likely be attributed to differences in ocean area, not body length.

**Fig. 2.**
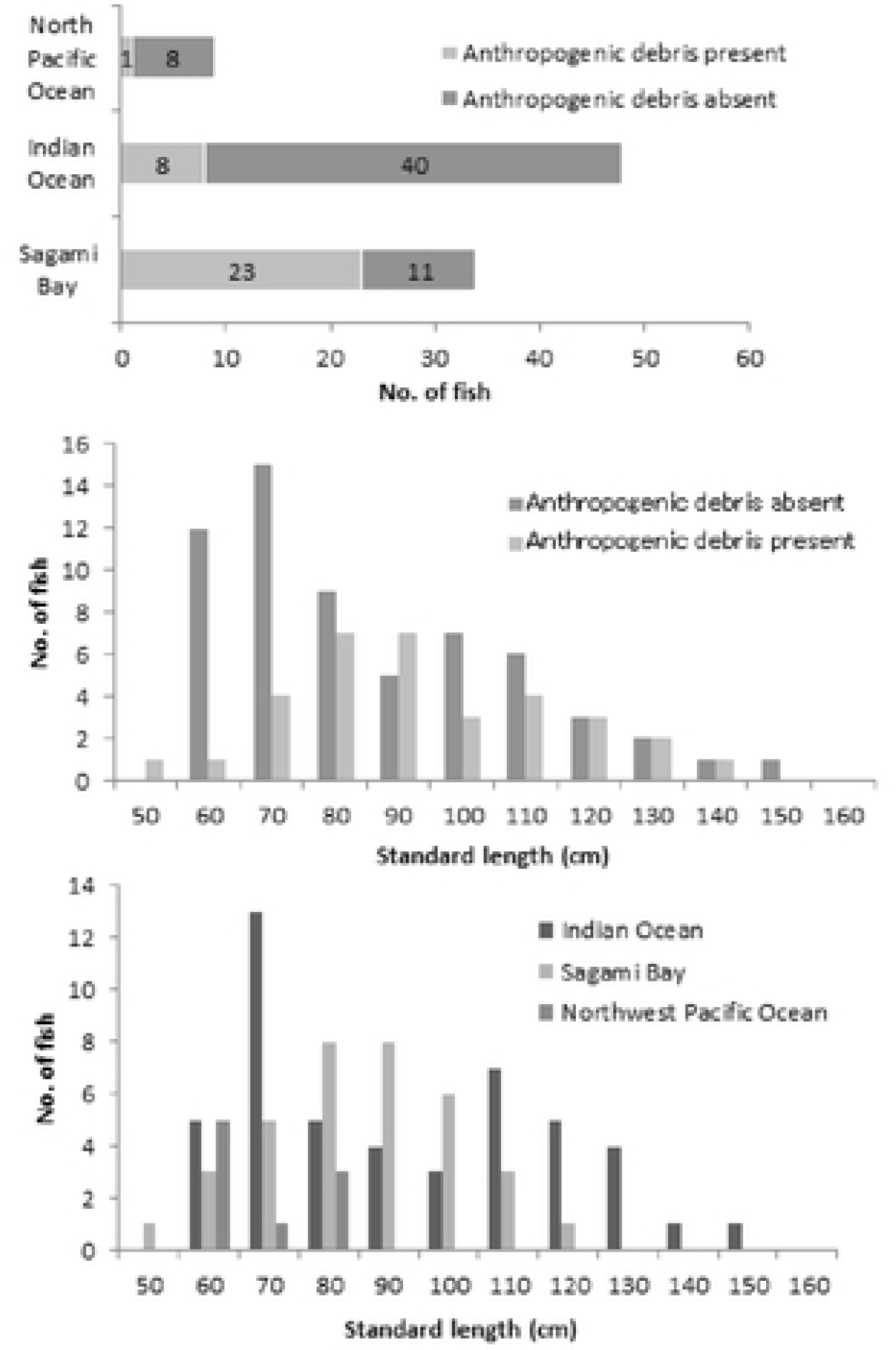
a. Incidence of anthropogenic debris ingestion by longnose lancetfish. b. Body length distribution of longnose lancetfih with and without ingested anthropogenic debris. c. Body length distribution of longnose lancetfish in different oceans.

### Relative proportions of anthropogenic debris and food items

The relationship between the number of anthropogenic and food items in the stomach contents is presented in Fig. 3. The figure shows a comparison of the number of both anthropogenic and natural items without considering their type or size. In some specimens, no items were found in the stomach; these included 5 of 34 specimens caught in Sagami Bay, 22 of 48 specimens caught in the Indian Ocean, and 2 of 9 specimens caught in the North Pacific Ocean. Data for these specimens were not included in the figure. According to Fig. 3, the two specimens in which anthropogenic items outnumbered natural items were those that were caught in Sagami Bay and Indian ocean. Because the residence time of anthropogenic debris and the time required to digest natural items in longnose lancetfish stomachs are not known, it is not possible to estimate the relative proportions of artificial debris and food organisms in the ocean. However, comparison of different ocean areas indicates that the proportion of anthropogenic items relative to natural items is substantially higher in Sagami Bay that in other ocean areas.

**Fig. 3.**
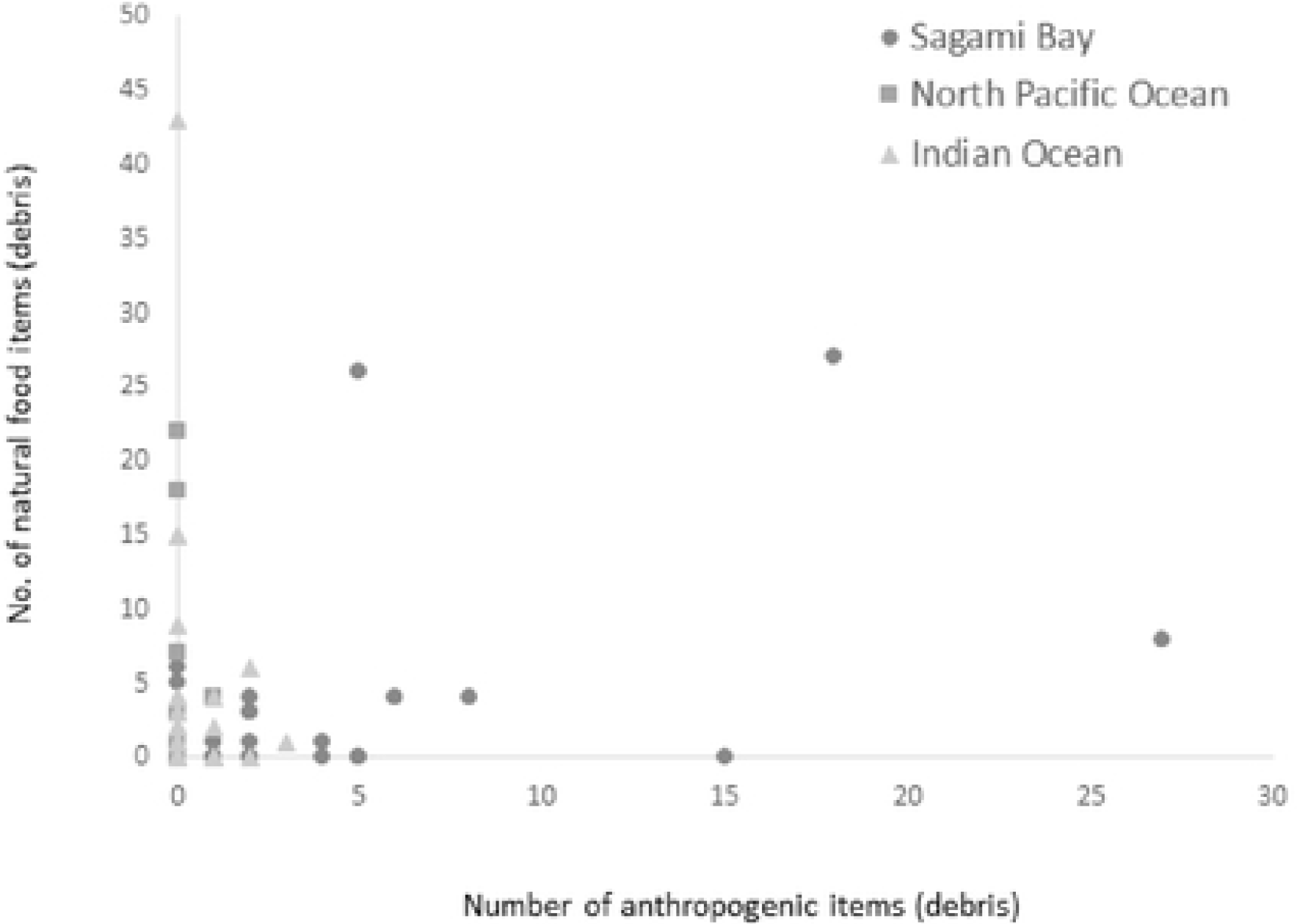
Relationship between number of ingested anthropogenic and natural items.

### Types of anthropogenic debris observed

The types of anthropogenic items recorded in longnose lancetfish stomachs are shown in Fig. 4. Of the 130 anthropogenic items observed, 117 were recorded in specimens caught in Sagami Bay, 12 items were recorded in specimens caught in the Indian Ocean, and one item was recorded in a specimen caught in the North Pacific Ocean. With the exception of one paper scrap, all of the items were products derived from petroleum. The most common items, accounting for over 50% of the items found, were sheet fragments. When bag fragments and food packing fragments believed to be plastic sheet fragments are also included, plastic sheet fragments account for more than 70% of the anthropogenic items recorded.

**Fig. 4.**
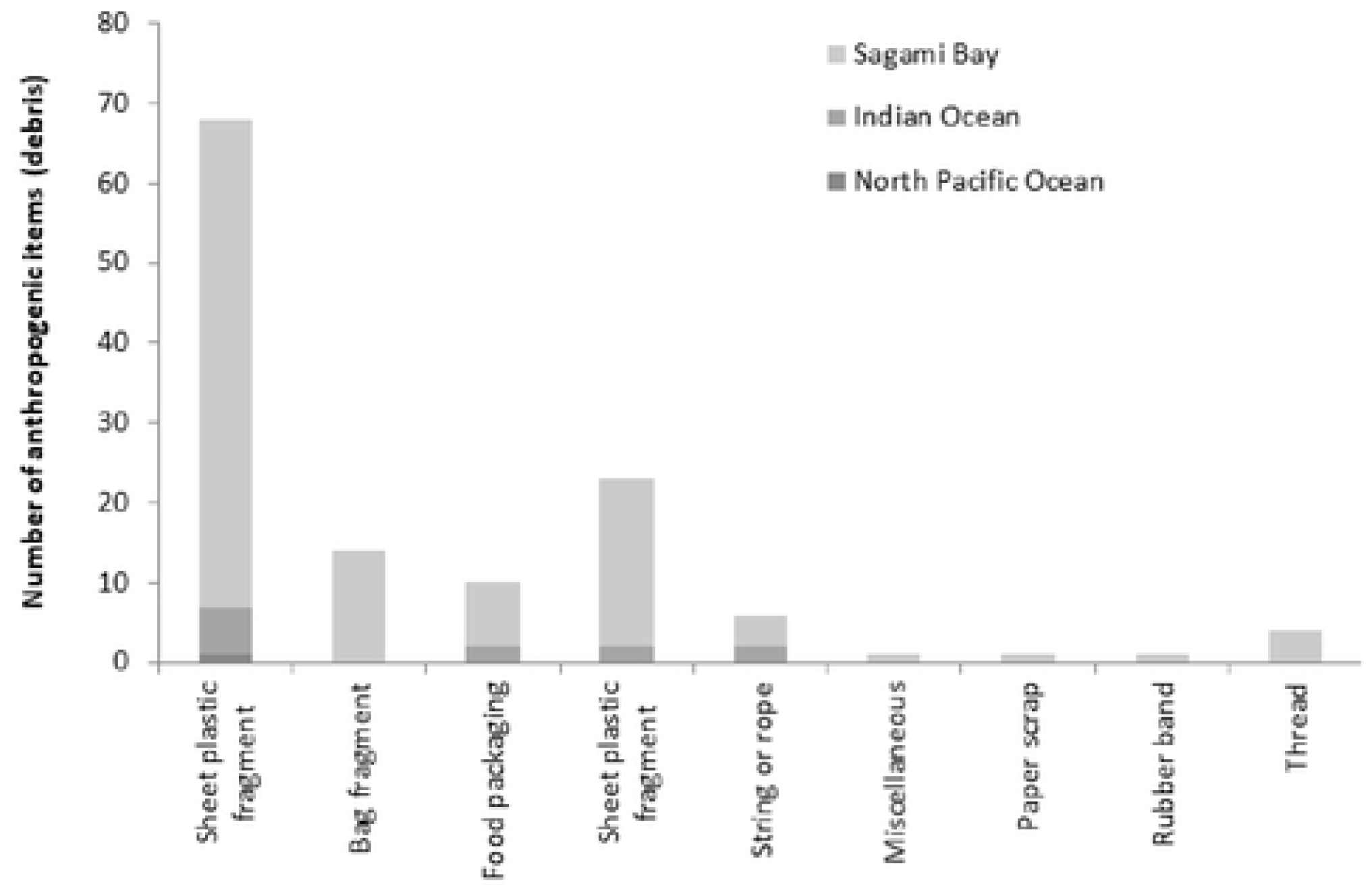
Number of ingested anthropogenic items by item type

**Fig. 5.**
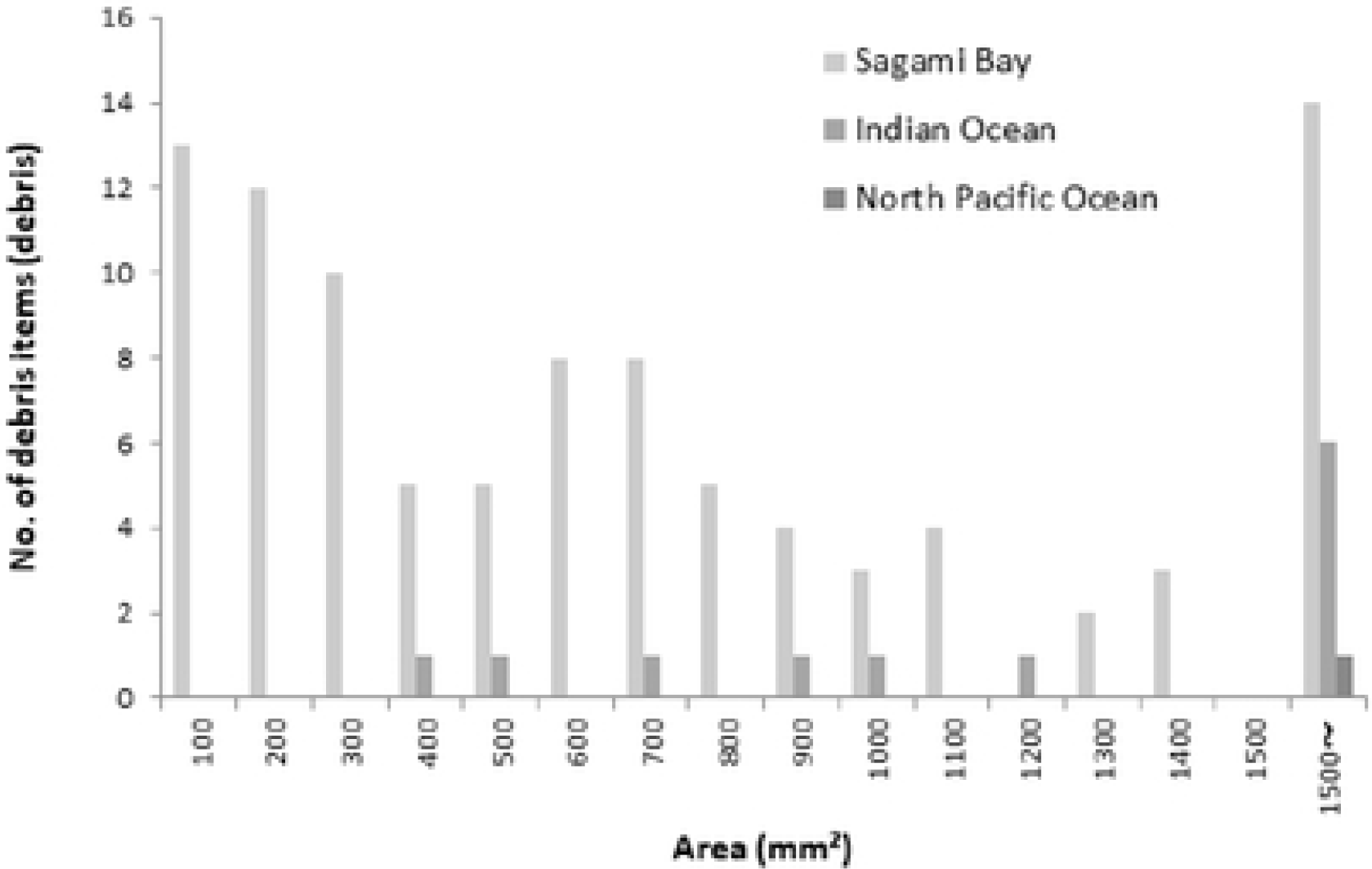
Frequency distribution of ingested anthropogeninc items of different sizes (areas)

### Relationship between size of anthropogenic debris and body length

Using photographs of the stomach contents of 25 specimens, we calculated the projected area of the anthropogenic items and investigated the relationship between particle area and body length for the specimens in which the debris was found. The smallest and largest anthropogenic items in the photographs measured 11 mm2 and 18,196 mm2, respectively. More than 80% of the items were less than 1,500 mm2, with the most common size ranging less than 100 mm2 (Fig. 6). No correlation was observed between body length and size of the anthropogenic items observed (Fig. 6).

**Fig. 6.**
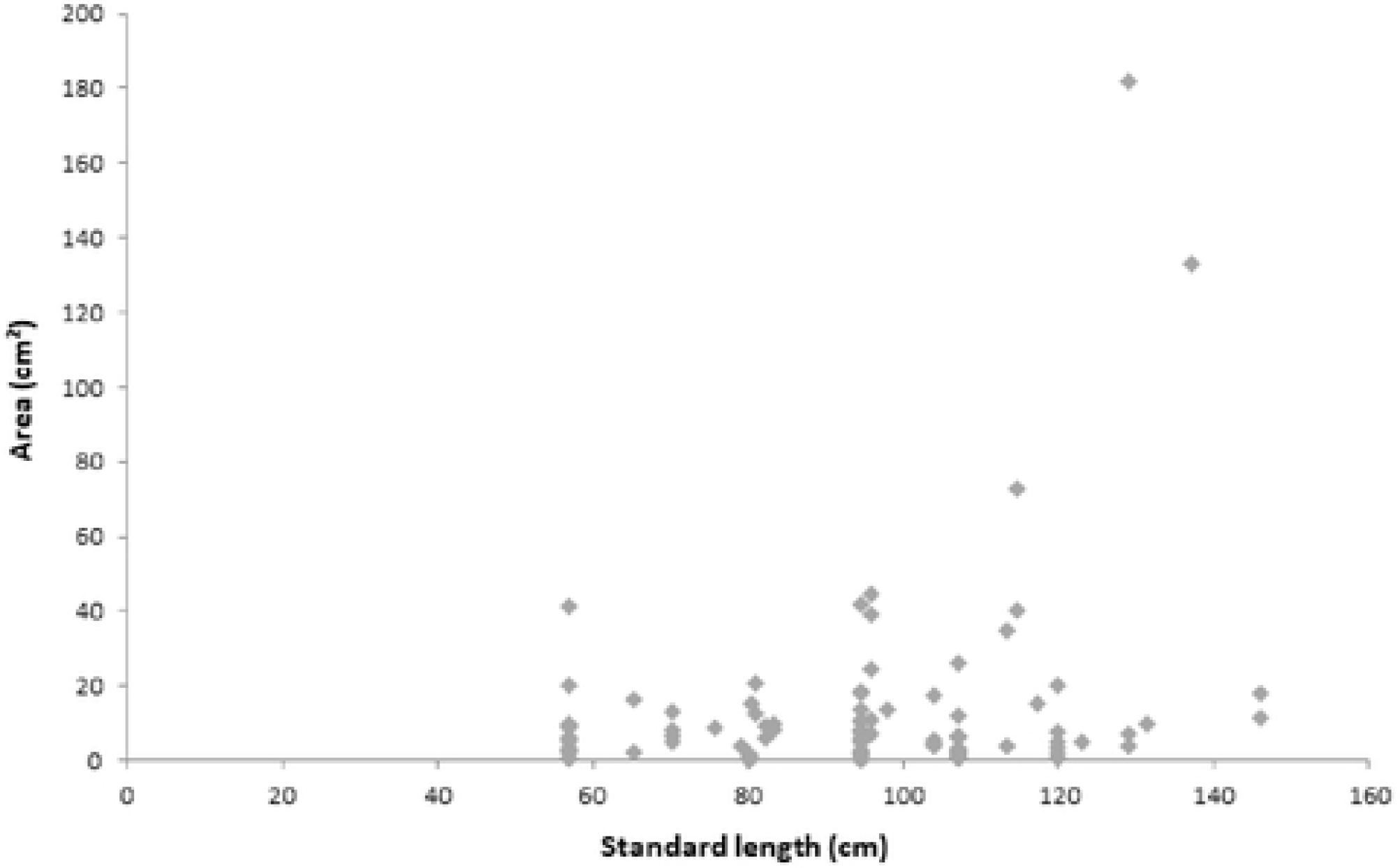
Relationship between longnose lancet fish standard length (SL) and size (surface area) of ingested anthropogenic items

### Material composition of ingested anthropogenic debris

The material composition of anthropogenic items collected from 19 specimens is shown in Fig. 7. Of the 71 fragments examined, 37 were composed of polypropylene (PP), 26 were polyethylene (PE), 4 was polycarbonate (PC), one was polyvinylidene chloride (PVDC), and 3was another plastic. PP and PE, whose specific gravities are lower than that of seawater, accounted for more than 90% of the observed items.

**FIG. 7.**
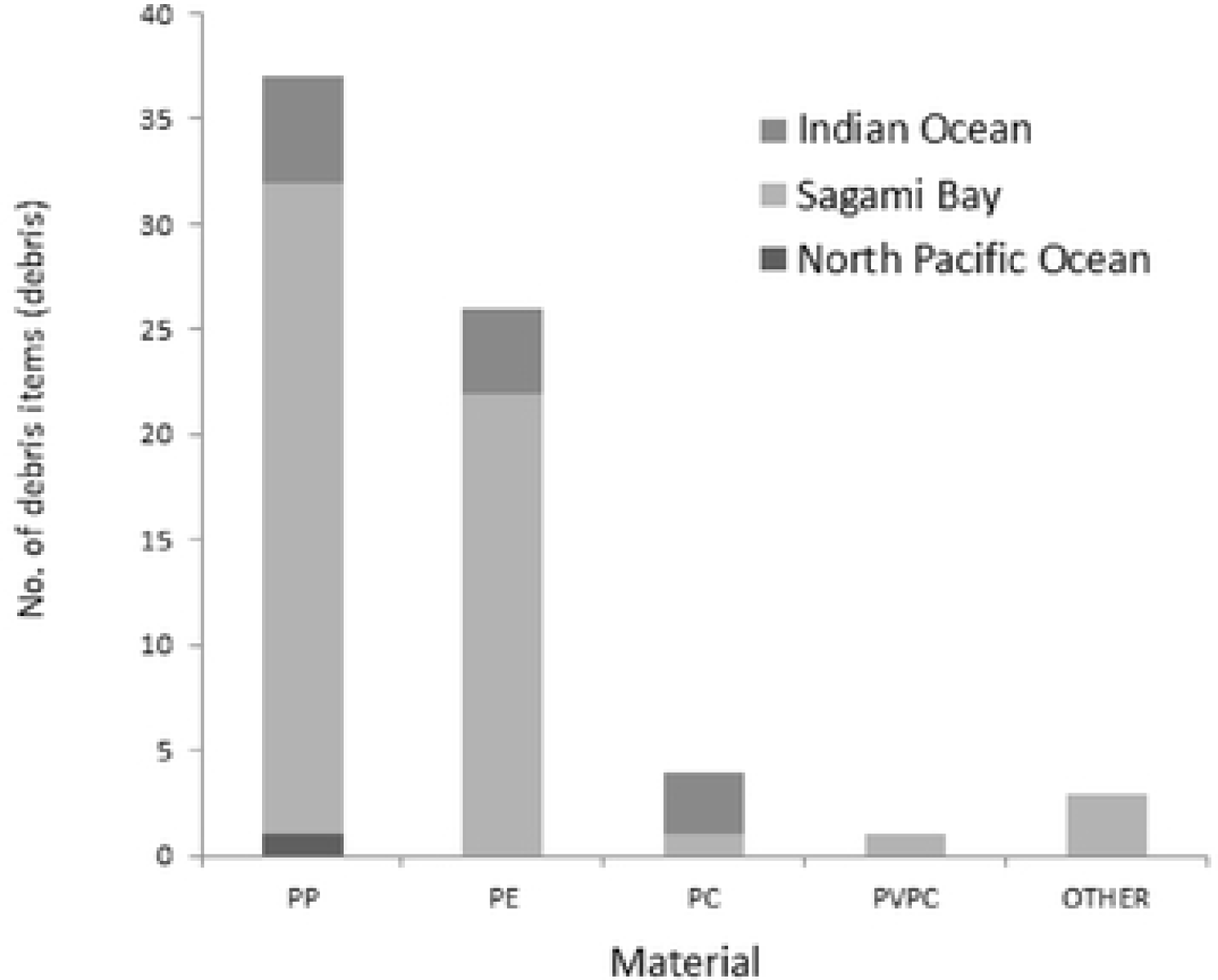
Number of ingested anthropogenic items by material composition

### Incidence of anthropogenic debris and capture depth

The estimated depths at which the different longnose lancetfish specimens were caught are presented in Fig. 8. Specimens were caught at a maximum depth of 306 m and a minimum depth of 36 m. Overall, most of the Sagami Bay specimens were caught at above 100 m, while the largest number of Indian Ocean specimens were caught at depths between 100 and 200 m, and the largest number of North Pacific Ocean specimens were caught below 150 m. No correlation was observed between body length and capture depth.

**Fig. 8.**
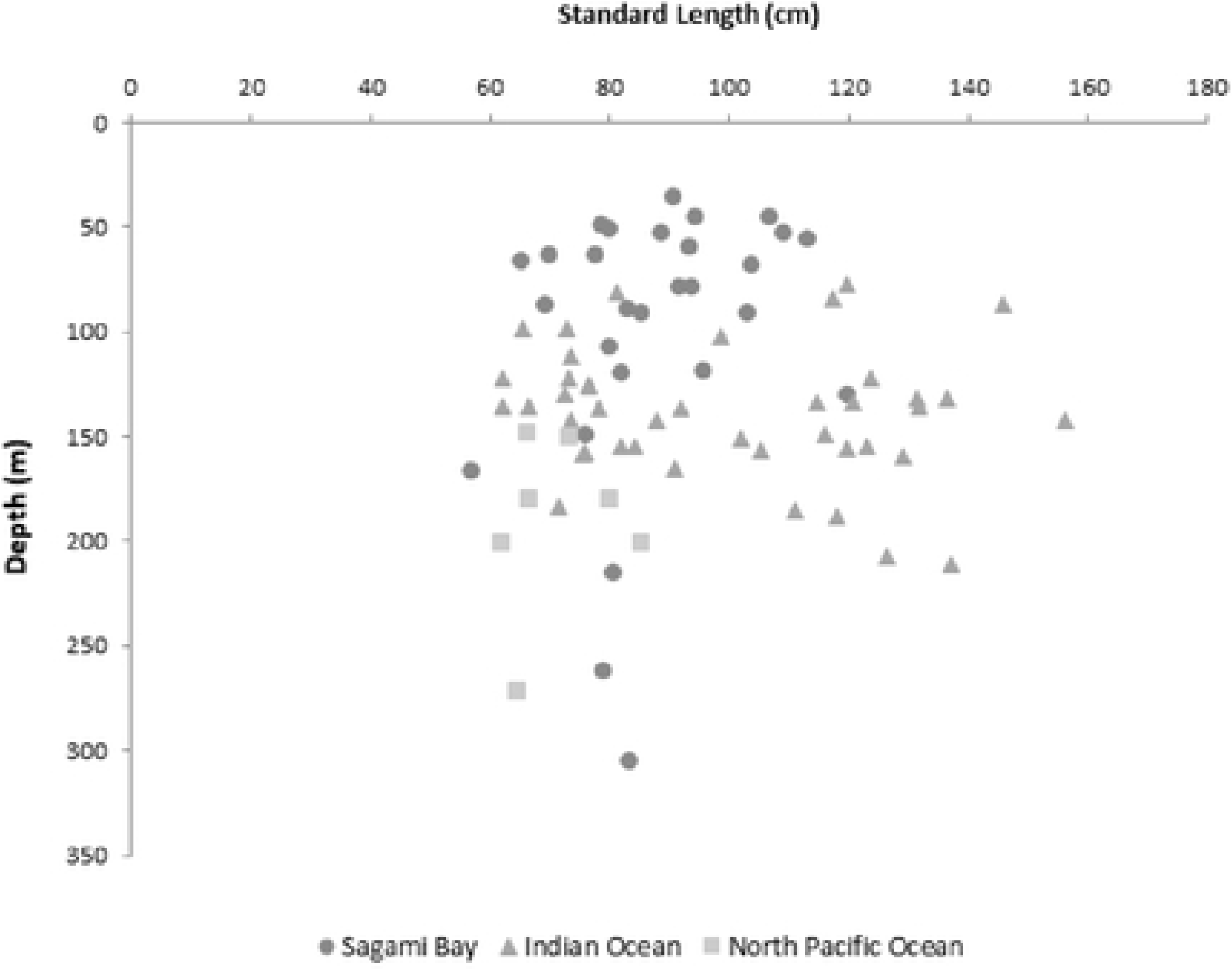
Relationship between longnose lancetfish total length and capture depth

We also examined the relationship between depth and stomach contents containing anthropogenic debris. The relationship between the number of anthropogenic items observed and capture depth is shown in Fig. 9. There were three specimens in which at least 10 anthropogenic items were found; these included two specimens caught at depths above 50 m and one specimen caught below 150 m. The greatest capture depth for any specimen with anthropogenic debris in its stomach (2 fragments) was 216 m. Although no correlation was observed between the number of fragments found and the capture depth.

**Fig. 9.**
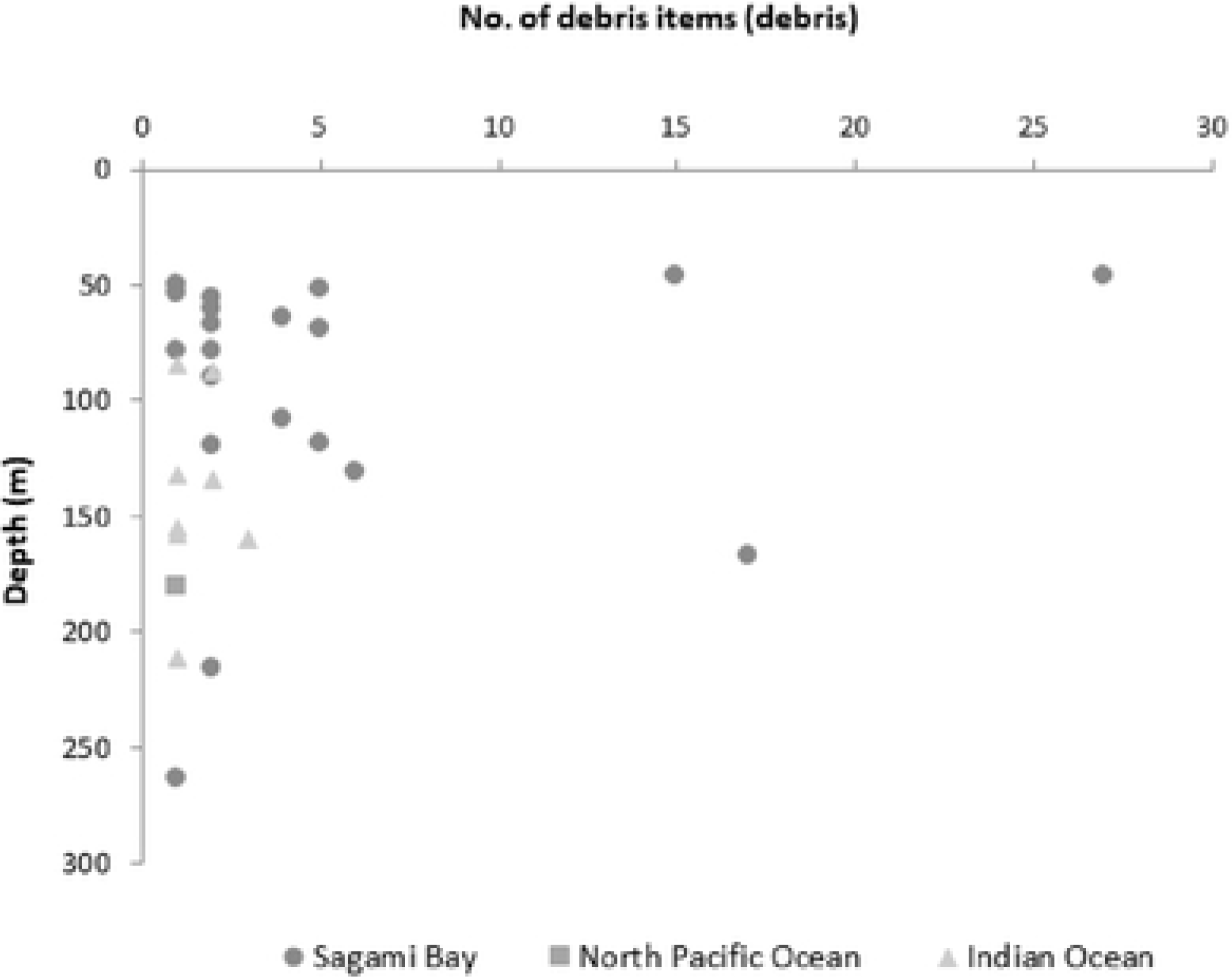
Relationship between number of ingested anthropogenic items and capture depth. Inset photograph: anthropogenic debris from the stomach of a longnose lancetfish caught at a depth of 216 m.

## Discussion

The feeding strategy of the longnose lancetfish is to eat everything in its path. If a longnose lancetfish senses an object in its environs, it will swallow it if it will fit in its mouth. The largest specimen in this study, which had a standard length of 131.8 cm, was found to have cannibalized a smaller longnose lancetfish whose standard length was 66.4 cm. It is considered that, if larger fragments of marine debris are suspended in the ocean, that they would be ingested by longnose lancetfish. Meanwhile, small fish measuring approximately 2 cm in length were found in the stomach of another specimen that had a standard length of 120 cm. Combined with the absence of a correlation between the size of anthropogenic items and standard length, these findings suggest that longnose lancetfish, regardless of body length, do not have a preference for prey of any size. This tendency is consistent with that reported by Jantz et al (2013)[19]. Although the longnose lancetfish has this feeding habit, the majority of anthropogenic items found in this study were 1500 mm2 or less in size. The anthropogenic debris in the stomach of the specimen caught at the greatest depth (263 m) consisted of a plastic sheet (373 mm2). This plastic sheet was PP (Fig. 9). More than 70% of the anthropogenic items found in this study were classified as plastic sheeting. Stomach content analysis revealed that more than 90% of the plastic fragments were composed of PP and PE, which have specific gravities that are less than that of seawater. Given that PP and PE plastic debris should therefore float on the water surface, the abundance of these plastics in the stomachs of the lancetfish examined suggests that these small fragments lost their buoyancy due to biofouling [23] and were either in the process of settling or were suspended in the water column as they are neutrally buoyant. In this study, none of the stomachs examined contained Styrofoam which are usually observed on sea level pop-ups. That is, it is suggested that longnose lancetfish do not ingest debris floating on the ocean surface but, rather, debris suspended in the epipelagic zone. Meanwhile, in the North Pacific Ocean survey, plastic fragments, fishing nets, and different types of rope accounted for 90% of ingested anthropogenic items. The composition of these materials differed substantially from materials found elsewhere, suggesting that the composition of anthropogenic debris suspended in the epipelagic zone differs by ocean region. Further, we think that these debris comprise small plastic items or sheet-like fragments resulting from the breakdown of larger objects that have lost their buoyancy and are suspended at depths between 0 and 200 m below the surface. Although the presence of vast amounts of plastic in the open ocean, recent studies show that its measured abundance is much smaller than expected [24]. There is the theory that a large part of the plastic has been degraded by either physical and biotic processes [25]. Our result showed that plastic debris are drifting from the middle layer to the deep layer as much as longnose lancetfish is mistaken for bait. The incidence of anthropogenic debris ingestion in different ocean areas ranged from 68% in Sagami Bay to 11% in the North Pacific Ocean, and 17% in the Indian Ocean. Comparing these results to previous studies, anthropogenic debris was found in approximately 70% of longnose lancetfish caught in Suruga Bay, and in 24% of longnose lancetfish caught near the North Pacific subtropical frontal zone [14],[19]. In our study, the result for Sagami Bay is similar to that for the catch in Suruga Bay, suggesting that there is an increased probability of encountering anthropogenic debris near human living environments. The incidence of anthropogenic debris ingestion has been shown to increase near areas such as the North Pacific subtropical frontal zone, which is known to accumulate marine debris [12]. Meanwhile, marine debris derived from plastic products was also found in longnose lancetfish caught in the Indian Ocean and the North Pacific Ocean, which are not frontal zones. The incidence of anthropogenic debris ingestion in the Indian Ocean was 17%, which was 11% in the Pacific Ocean. As a result of statistics analyzes, there was no significant difference between the two values. From these facts, there is a possibility that there may be high density areas in the Indian Ocean that are equal to or higher than the Pacific Ocean.

In addition, microplastics (< 5mm), which are smaller than the debris investigated in this study, have been found in the remotest of locations, namely the Antarctic Ocean [26]. Similarly, microfibers have been detected in arthropods and cnidaria inhabiting the deep ocean floor at depths of 1,062 m and 1,783 m in the southwestern Indian Ocean and the central Atlantic Ocean, respectively [27]. The results of our study suggest that marine debris not only exists at the ocean surface and on the ocean floor, but also throughout the epipelagic zone in the form of plastic sheet fragments that have started to degrade; this plastic debris is becoming widely distributed in the Pacific and Indian Oceans. We conclude that ocean areas uncontaminated by marine debris are beginning to disappear. At present, we do not know how long longnose lancetfish swim, how large their ranges to feed area, how long it takes for ingested debris to be excreted, or even if it is excreted at all. For this reason, we are not able to estimate the specific concentration of marine debris in the epipelagic zone from longnose lancetfish. However, relative comparisons of different ocean areas can be made by comparing the stomach contents of longnose lancetfish. We believe that comparison of a wider range of ocean areas and continuous monitoring of relative amounts of marine debris in the epipelagic zone are necessary.

## Acknowledgements

We gratefully acknowledge for collecting our sample to the captains and crews of the T/V Umitaka-Maru, T/V Shinyo-Maru and T/V Seiyo-maru. We want to thank past and present members of our laboratory for their help. Part of this research was supported by the JSPS KAKENHI Grant Number 16H017

